# *De novo* transcriptome assembly for the spiny mouse (*Acomys cahirinus*)

**DOI:** 10.1101/076067

**Authors:** Jared Mamrot, Roxane Legaie, Stacey J Ellery, Trevor Wilson, David K. Gardner, David W. Walker, Peter Temple-Smith, Anthony T. Papenfuss, Hayley Dickinson

## Abstract

Background: Spiny mice of the genus *Acomys* are small desert-dwelling rodents that display physiological characteristics not typically found in rodents. Recent investigations have reported a menstrual cycle and scar free-wound healing in this species; characteristics that are exceedingly rare in mammals, and of considerable interest to the scientific community. These unique physiological traits, and the potential for spiny mice to accurately model human diseases, are driving increased use of this genus in biomedical research. However, little genetic information is currently available for *Acomys*, limiting the application of some modern investigative techniques. This project aimed to generate a reference transcriptome assembly for the common spiny mouse (*Acomys cahirinus*).

Results: Illumina RNA sequencing of male and female spiny mice produced 451 million, 150bp paired-end reads from 15 organ types. An extensive survey of *de novo* transcriptome assembly approaches of high-quality reads using Trinity, SOAPdenovo-Trans, and Velvet/Oases at multiple kmer lengths was conducted with 49 single-kmer assemblies generated from this dataset, with and without *in silico* normalization and probabilistic error correction. Merging transcripts from 49 individual single-kmer assemblies into a single meta-assembly of non-redundant transcripts using the EvidentialGene ‘*tr2aacds*’ pipeline produced the highest quality transcriptome assembly, comprised of 880,080 contigs, of which 189,925 transcripts were annotated using the SwissProt/Uniprot database.

Conclusions: This study provides the first detailed characterization of the spiny mouse transcriptome. It validates the application of the EvidentialGene ‘*tr2aacds*’ pipeline to generate a high-quality reference transcriptome assembly in a mammalian species, and provides a valuable scientific resource for further investigation into the unique physiological characteristics inherent in the genus *Acomys*.

## Background

The Common or Cairo spiny mouse (*Acomys cahirinus*) is a small rodent species endemic to the semi-arid deserts of Africa and the Middle East (Wilson & Reeder, 2005). Used in research to model human disease, spiny mice exhibit physiological characteristics not typically found in rodents: they exhibit a precocial pattern of development (Brunjes, 1990; Dickinson et al., 2005), atypical synthesis of hormones such as cortisol and dehydroepiandosterone (Lamers, 1986; Quinn et al., 2013; 2015), and a menstrual cycle (Bellofiore et al., 2016). These traits are common to humans and other higher order primates, but rare in other mammals. For example, menstruation has been identified in only six non-primate species, none of which are rodents (Emera et al., 2012). The discovery of human-like physiological characteristics in a rodent is highly valuable for those in the scientific community looking to model human conditions. The true value of the common spiny mouse as an animal model has yet to be fully realized, as many fundamental aspects of their biology are still to be explored; for instance, there is little genetic information available for this species.

Publically available genetic information for the spiny mouse consists of the mitochondrial genome (Hadid et al., 2014), and two RNA sequencing (RNA-Seq) datasets: PRJNA184055 (Fushan et al., 2015), and PRJNA292021 (Gawriluk et al., 2016). These next-generation sequencing (NGS) datasets were created with specific aims: to establish incipient sympatric speciation as a mode of natural selection in mammals inhabiting divergent microclimates (Hadid et al., 2014), to examine the molecular basis for natural variation in mammalian lifespan (Fushan et al., 2015), and to characterize and investigate another characteristic unique to *Acomys*: scar-free wound healing and skin regeneration (Gawriluk et al., 2016). *De novo* assembly of NGS reads was conducted for each specific organ/tissue sequenced in these projects, however the accuracy and completeness of the resulting assemblies were not explicitly described. Accurate identification of differentially expressed genes is dependent on accurate read mapping (Garber et al., 2011). The use of a high-quality species-specific reference sequence containing transcripts from multiple organ types promotes accurate mapping, thereby facilitating accurate gene expression analysis.

RNA-Seq provides an unprecedented opportunity for cost-effective, large-scale genetic analysis in non-model organisms for which a genome sequence is unavailable. *De novo* assembly of millions/billions of RNA-Seq reads into a reference transcriptome can provide a valuable scientific resource, with applications in phylogenetic analysis (Robertson & Cornman, 2014), novel gene identification (Maudhoo et al., 2015), RNA editing (Athanasiadis et al., 2004) and alternative splicing investigation (Du et al., 2015), qPCR primer design (Zieliński et al., 2014), development and refinement of bioinformatics software (Bens et al., 2016), augmenting proteomic research (Francischetti et al., 2013), and investigation of gene expression profiles underlying complex physiological traits (Carneiro et al., 2014, Shimoyama et al., 2016). Surprisingly, there is no universal protocol for the assembly process. Despite the availability of protocols for *de novo* transcriptome assembly using popular software packages (eg. Haas et al., 2013), and practical guidelines such as the comprehensive Oyster River protocol (MacManes, 2016; http://oyster-river-protocol.rtfd.org/), the assembly pipeline commonly requires considerable optimisation to obtain high quality, meaningful results from each dataset (Robertson et al., 2010; Surget-Groba & Montoya-Burgos, 2010).

Here, we describe the application of a variety of *de novo* transcriptome assembly methods, utilizing both single- and multi-kmer approaches, with unique transcripts from multiple assemblies combined to address our primary aim: the generation of a comprehensive *de novo* transcriptome assembly for the common spiny mouse (*Acomys cahirinus*).

## Methods

### Sample preparation and sequencing

Tissues were collected from male and female adult spiny mice, and placenta from three pregnant females. Total RNA was extracted using Trizol reagent from skin, lung, liver, small intestine, kidney, adrenal gland, brain, thymus, spleen, diaphragm, heart, skeletal muscle, testis, ovary, and placenta. Quantitation and quality control was performed using Agilent Bioanalyzer and Qubit, with all samples returning RIN scores >7.0. The Illumina TruSeq Stranded Total RNA Library Prep kit (Illumina, Hayward, CA) was used to generate indexed cDNA libraries for each RNA sample. These were sequenced to produce paired-end 150bp reads on the Illumina HiSeq1500. Image analysis, raw nucleotide base calling, conversion from .bcl to .fastq format, and quality filtering were conducted using Illumina CASAVA v1.8.2 software (Illumina, Hayward, CA). Data yield (in Mbases), percent pass-filter (%PF), read count, raw cluster percentage per lane, and quality (%Q30) were examined using the Illumina Sequencing Analysis Viewer 1.8.46 (Illumina, Hayward, CA).

### Data processing

#### Sequence reads were quality checked using FastQC v0.11.3

(http://www.bioinformatics.babraham.ac.uk/projects/fastqc/). Adapter sequences and low quality bases (Q<20) were trimmed from 3’ ends using trim-galore (ver: 0.4.0; http://www.bioinformatics.babraham.ac.uk/projects/trim_galore/), which wraps cutadapt v0.9.5 (Martin, 2011). Reads with average quality scores lower than 20 and read pairs in which either forward or reverse reads were trimmed to fewer than 35 nucleotides were discarded. Remaining reads were assessed again using FastQC.

Filtering of poor quality reads was conducted using Trimmomatic v0.30 (Bolger et al., 2013) with settings ‘LEADING:3 TRAILING:3 SLIDINGWINDOW:4:20 AVGQUAL:20 MINLEN:35’. Up to three nucleotides with quality scores lower than 20 were trimmed from the 3’ and 5’ read ends. Reads with an average quality score lower than 20, and reads with a total length of fewer than 35 nucleotides were removed.

Probabilistic error correction was performed using SEECER (Hai-Son et al., 2013) with default parameters, remaining high-quality reads were again examined with FastQC. Both error-corrected and non-error-corrected reads were subjected to *de novo* assembly.

#### *De novo* transcriptome assembly

Reads were assembled using either SOAPdenovo-Trans v1.03 (Xie et al., 2014), Trinity package r20140413p1 (Grabherr et al., 2011; available at http://trinityrnaseq.github.io), or Velvet v1.2.10 (Zerbino & Birney, 2008) in combination with Oases v0.2.08 (Schultz et al., 2012) with default parameters, except where indicated. The single-kmer assemblies were performed with and without digital normalization and probabilistic error correction as described in Figure 1. Reads were subjected to digital normalization using the script bundled with the Trinity package, or with BBNorm (part of the BBMap v35.85 package; Bushnell, 2016: *http://sourceforge.net/projects/bbmap*) (Crusoe et al., 2015).

SOAPdenovo-Trans parameters: “max_rd_len=150, rd_len_cutof=150, avg_ins=192, reverse_seq=0, asm_flags=3” with kmer lengths: 21, 23, 25, 27, 29, 31, 35, 41, 51, 61, 71, 81, 91. Trinity was used at kmer length 25, with parameters: “--normalize_reads --seqType fq --JM 100G --CPU 20 --min_kmer_cov 2”. Reads were assembled with Velvet at kmer lengths 21, 23, 25, 27, 29, 31, 35, 41, 51, 61, 71, 81, 91, 101, 111, 121. Velvet was compiled with parameters “MAXKMERLENGTH=141 BIGASSEMBLY=1 LONGSEQUENCES=1 OPENMP=1”. Velveth was run using “25,37,2 -shortPaired -fastq -separate” and “41,131,10 -shortPaired -fastq -separate”. Insert lengths of the fragments were estimated with CollectInsertSizeMetrics in Picard Tools version 1.90 (http://broadinstitute.github.io/picard/). Velvetg was run with parameters “-read_trkg yes -ins_length 215”. Oases was run with parameters “-min_trans_lgth 100 -ins_length 215”.

Assembly statistics were computed using the TrinityStats.pl from the Trinity package, and log files produced by SOAPdenovo-Trans and Velvet/Oases.

#### Collating non-redundant transcripts from multiple assemblies

The *tr2aacds* pipeline from the EvidentialGene package (Gilbert, 2013: http://arthropods.eugenes.org/EvidentialGene/about/EvidentialGene_trassembly) was used to identify and collate non-redundant transcripts from each individual transcriptome assembly. The *tr2aacds* pipeline predicts amino acid sequences and transcript coding sequences, removes transcript redundancy based on coding potential, removes sequence fragments, clusters highly similar sequences together into loci, and classifies non-redundant transcripts as ‘primary’ or ‘alternative’. Transcripts that scored poorly were removed, with remaining ‘primary’ and ‘alternative’ transcripts from each single-kmer assembly collated into a final ‘clustered assembly’ (CA).

#### Assessing accuracy and completeness of each assembly

Accuracy and completeness was assessed in all assemblies (single-kmer and CA) using BUSCO v1.1b1 (Simão et al., 2015) to establish the presence or absence of universal single copy orthologs common to vertebrates and eukaryotes. Accuracy was assessed by the proportion of original sequence reads mapped (‘backmapping’) to each assembly using Bowtie v0.12.9 (Langmead et al., 2009) with settings: ‘-q --phred33-quals -n 2 -e 99999999 -l 25 -I 1 -X 1000 -p 12 -a -m 200 --chunkmbs 256’. Independent RNA-Seq reads were obtained from the NCBI sequence read archive (SRA): datasets SRR636836, SRR636837, and SRR636838 were obtained from project PRJNA184055, and datasets SRR2146799 - SRR2146807 from project PRJNA292021. These reads were generated from liver (Fushan et al., 2015) and skin (Gawriluk et al., 2016) and neither organ was subjected to treatment i.e. these reads correspond to ‘control’ groups in their corresponding experiments. The independent RNA-Seq reads were aligned using Bowtie, with settings as specified above. The proportion of mapped reads was calculated using samtools flagstat with default parameters (Li et al., 2009). Structural integrity was examined using TransRate v1.0.2 (Smith-Unna et al., 2016) with default settings, in which Salmon v0.6.0 (Patro et al., 2015) and SNAP-aligner v0.15 (Zaharia et al., 2011) were implemented. Redundancy in assembled transcripts was assessed by the proportion of highly similar contiguous sequences (contigs), clustered using CD-HIT-EST v4.6.5 (Li & Godzik, 2006; Fu et al., 2012; Huang et al., 2010) with settings ‘-c 0.95 -n 8 -p 1 -g 1 -M 200000 -T 8 -d 40’. Further clustering at 90%, 95%, and 100% similarity was conducted on a representative single-kmer assembly to assess contig redundancy.

#### Annotation

The best performing assembly was annotated using the Trinotate pipeline (ver2.0.2, http://trinotate.github.io/). In brief, *de novo* transcripts were aligned against the SwissProt/Uniprot database (accessed 7th January 2016) using BLASTx and BLASTp (Altschul et al., 1990). Transdecoder v2.0.1 (https://transdecoder.github.io/) was used to predict ORFs, with BLASTp performed using predicted ORFs as the query and Swissprot/Uniprot database as the target. HMMER v3.1b1 and Pfam v27 databases (Finn, Clements & Eddy, 2011) were used to predict protein domains. SignalP v4.1 (Nielsen et al., 2011; Peterson et al., 2011) was used to predict signal peptides, RNAmmer v1.2 (Lagesen et al., 2007) to predict rRNA, and TMHMM v2.0 (Krogh et al., 2001) to predict transmembrane helices within the predicted ORFs. Annotations were loaded into an SQL database. GO terms linked to the SwissProt entry of the best BLAST hit were used for ontology annotation. GO functional classifications and plotting was performed by WEGO (http://wego.genomics.org.cn) (Ye et al., 2006).

## Results and Discussion

### Raw data

Raw reads are available from the NCBI SRA under project number PRJNA342864. In total, 451 million read pairs were generated, with yield, proportion aligned, error rate, intensity, and GC content provided in Table 1. Further summary statistics for data yield, percent pass-filter (%PF), raw cluster percentage per lane, and quality score summary are provided in Supplementary Table S1. Filtering of poor quality read pairs (Q<30) removed 32% of the original 451 million read pairs, with 305 million high-quality paired reads used for assembly. FastQC results for raw, filtered and *in silico* normalized data is provided in Supplementary Figure S1.

**Table 1.** Spiny mouse RNA-Seq statistics

|  | Forward | Reverse | Total |
| --- | --- | --- | --- |
| Yield total (G) | 45.9 | 45.9 | 91.77617 |
| Aligned (%) | 0.45 | 0.25 | 0.3509741 |
| Error rate (%) | 0.62 | 1.6 | 1.003602 |
| Intensity cycle 1 | 3255 | 3054 | 3154.379 |
| % >= Q30 | 80.5 | 46.5 | 63.47869 |
| Total raw reads |  |  | 611,841,080 |
| Total read pairs (Q>=30) |  |  | 305,920,540 |
| GC content |  |  | 45% |

### Transcriptome assembly

In total, 49 single-kmer transcriptome assemblies were produced from 305 million paired reads, with and without digital normalization and probabilistic error correction, as described in Figure 1. Detailed metrics for all transcriptome assemblies are provided in Supplementary Table S2. Digital normalization reduced the size of the dataset by >90%, however assemblies constructed using normalized data contained fewer BUSCO reference orthologues (Figure 2), had poorer backmapping rates (Figure 3), poorer mapping of independent reads (Figure 4), and poorer TransRate scores (Figure 5), compared to assemblies generated from non-normalized data.

**Figure 1.**
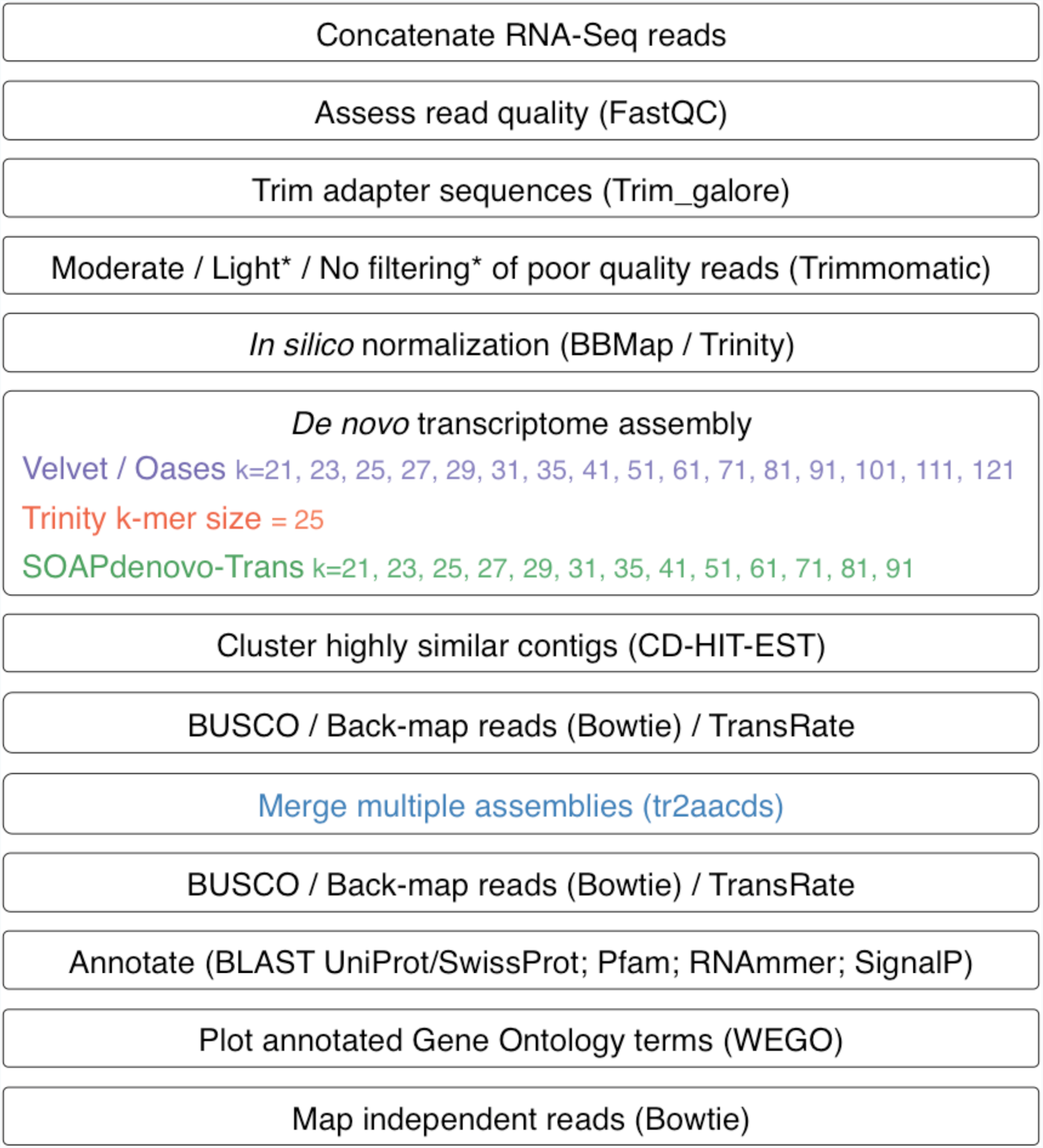
Flow chart of transcriptome assembly pipeline. *SEECER probabilistic error correction conducted on these datasets.

**Figure 2.**
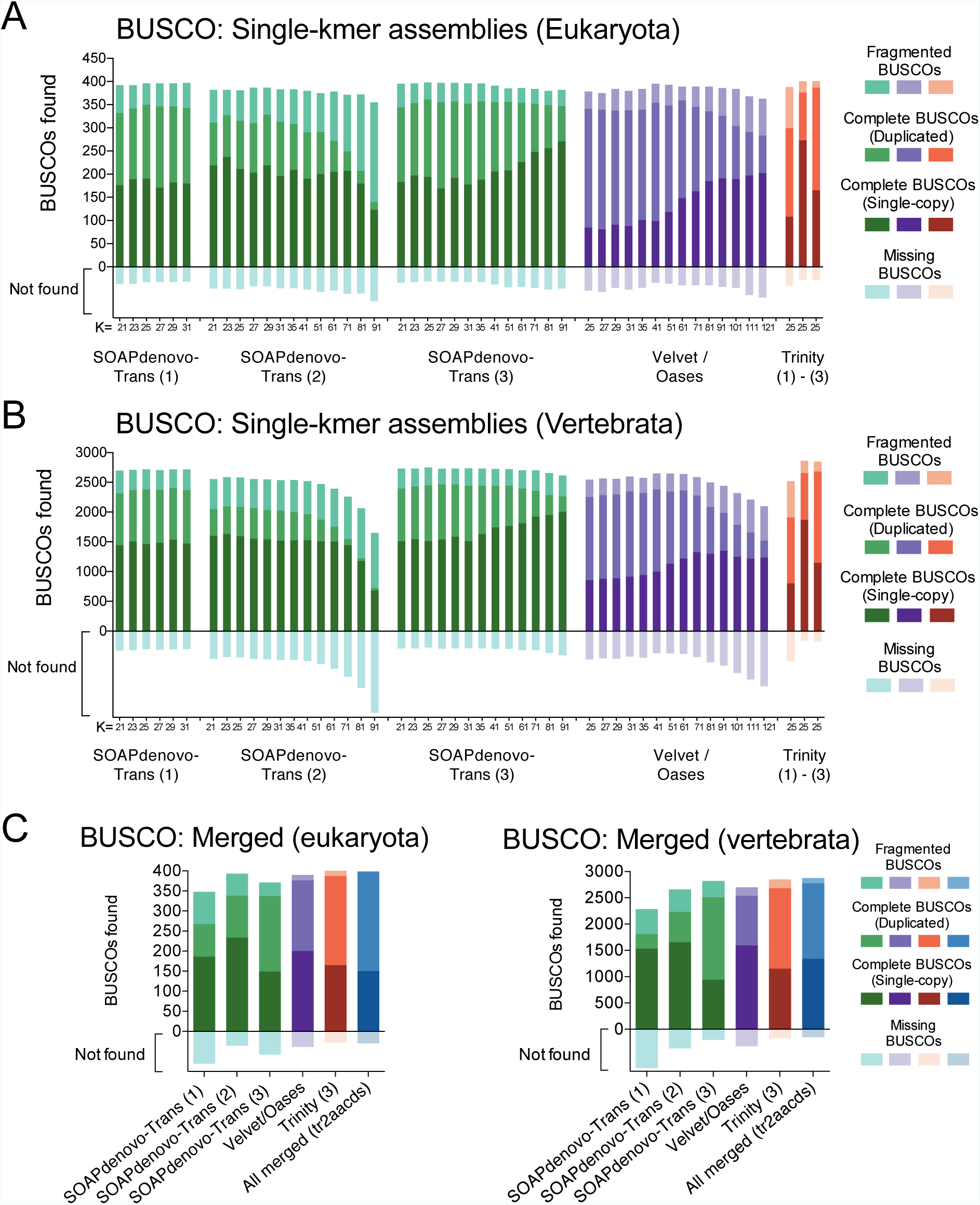
BUSCOs identified in each single-kmer transcriptome assembly. SOAPdenovo-Trans (1): no filtering; SOAPdenovo-Trans (2): light filtering of poor quality read pairs, combined with in silico normalization; SOAPdenovo-Trans (3): moderate filtering of poor quality read pairs; Velvet/Oases: moderate filtering of poor quality read pairs; Trinity (1): light filtering of poor quality read pairs, combined with in silico normalization; Trinity (2): light filtering of poor quality read pairs, combined with SEECER error correction; Trinity (3): moderate filtering of poor quality read pairs; CA: tr2aacds ‘clustered assembly’.

**Figure 3.**
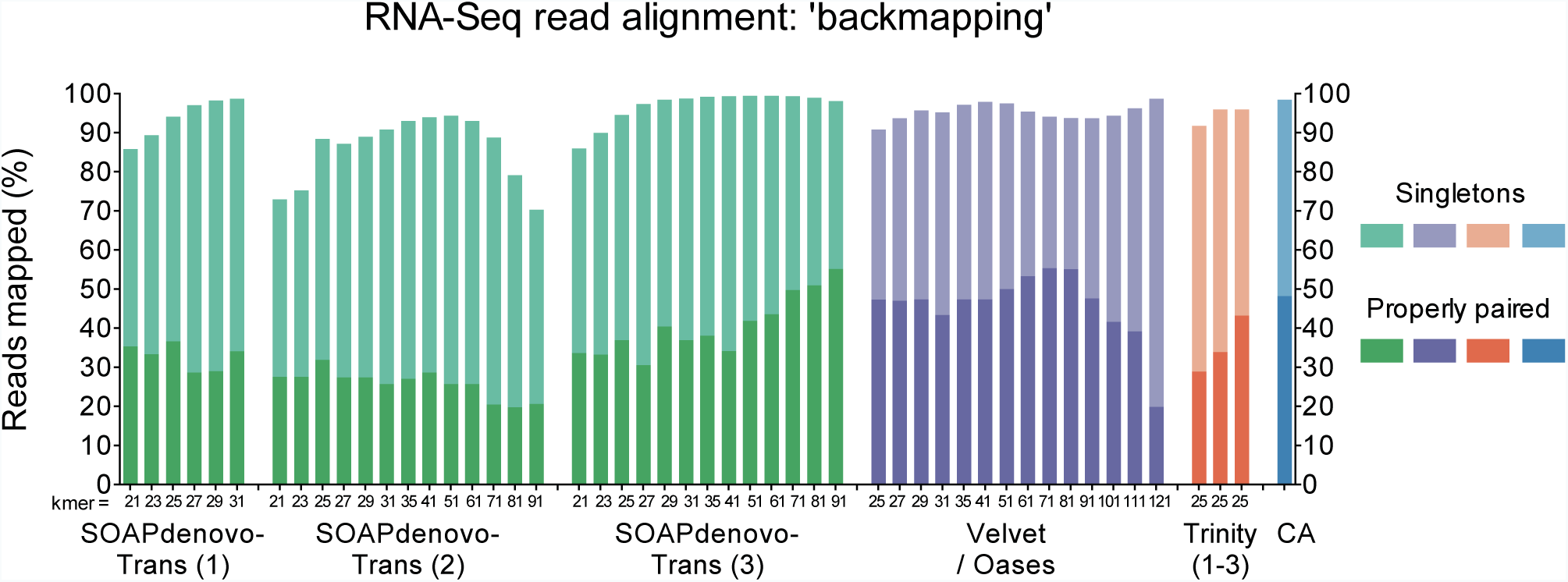
Proportion of reads backmapping to each transcriptome assembly. SOAPdenovo-Trans (1): no filtering; SOAPdenovo-Trans (2): light filtering of poor quality read pairs, combined with in silico normalization; SOAPdenovo-Trans (3): moderate filtering of poor quality read pairs; Velvet/Oases: moderate filtering of poor quality read pairs; Trinity (1): light filtering of poor quality read pairs, combined with in silico normalization; Trinity (2): light filtering of poor quality read pairs, combined with SEECER error correction; Trinity (3): moderate filtering of poor quality read pairs; CA: tr2aacds ‘clustered assembly’.

**Figure 4.**
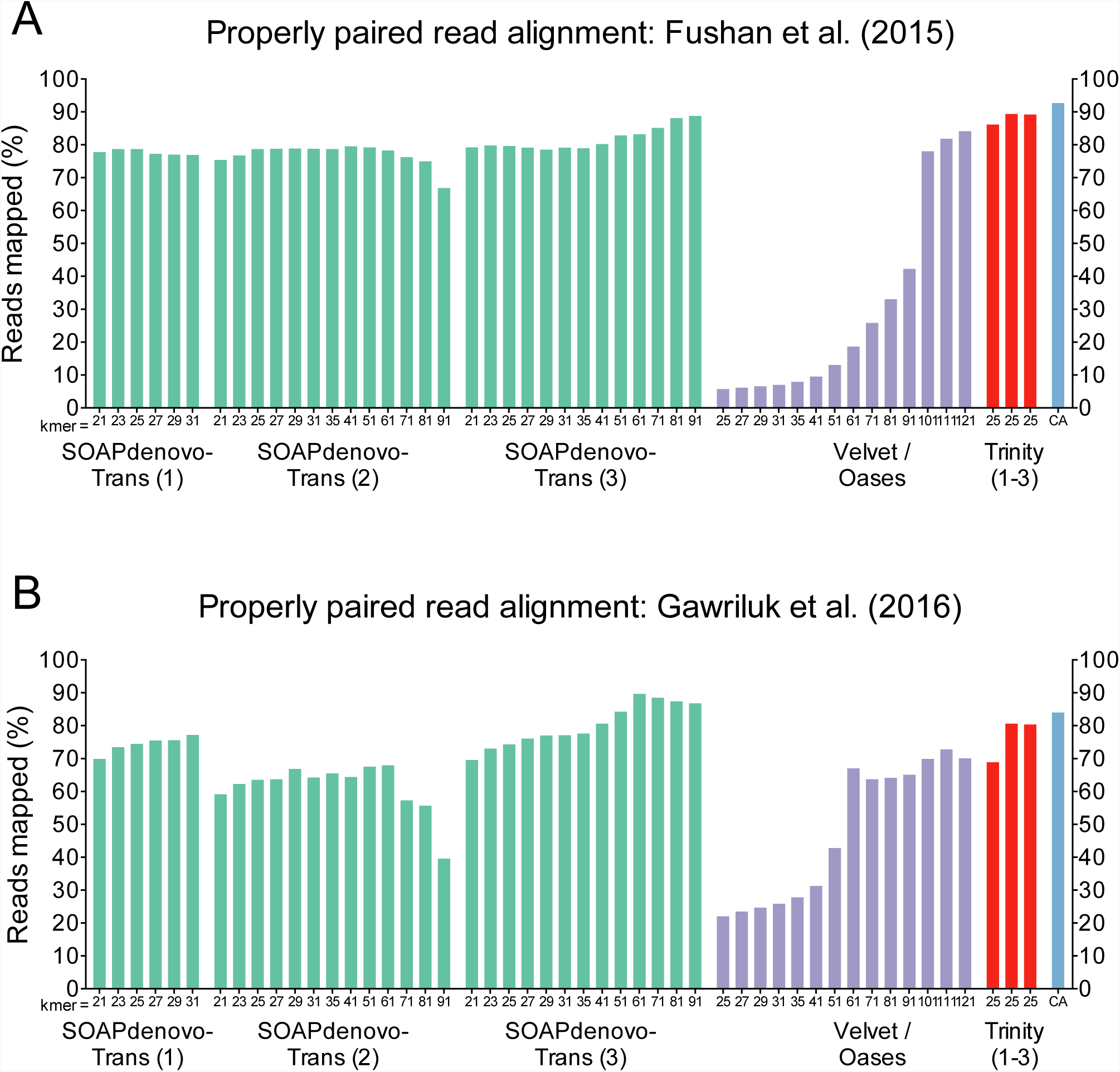
Proportion of independent reads mapping to each transcriptome assembly for (A) PRJNA184055 (Fushan et al., 2015), and (B) PRJNA300275 (Gawriluk et al., 2016). SOAPdenovo-Trans (1): no filtering; SOAPdenovo-Trans (2): light filtering of poor quality read pairs, combined with in silico normalization; SOAPdenovo-Trans (3): moderate filtering of poor quality read pairs; Velvet/Oases: moderate filtering of poor quality read pairs; Trinity (1): light filtering of poor quality read pairs, combined with in silico normalization; Trinity (2): light filtering of poor quality read pairs, combined with SEECER error correction; Trinity (3): moderate filtering of poor quality read pairs; CA: tr2aacds ‘clustered assembly’.

**Figure 5.**
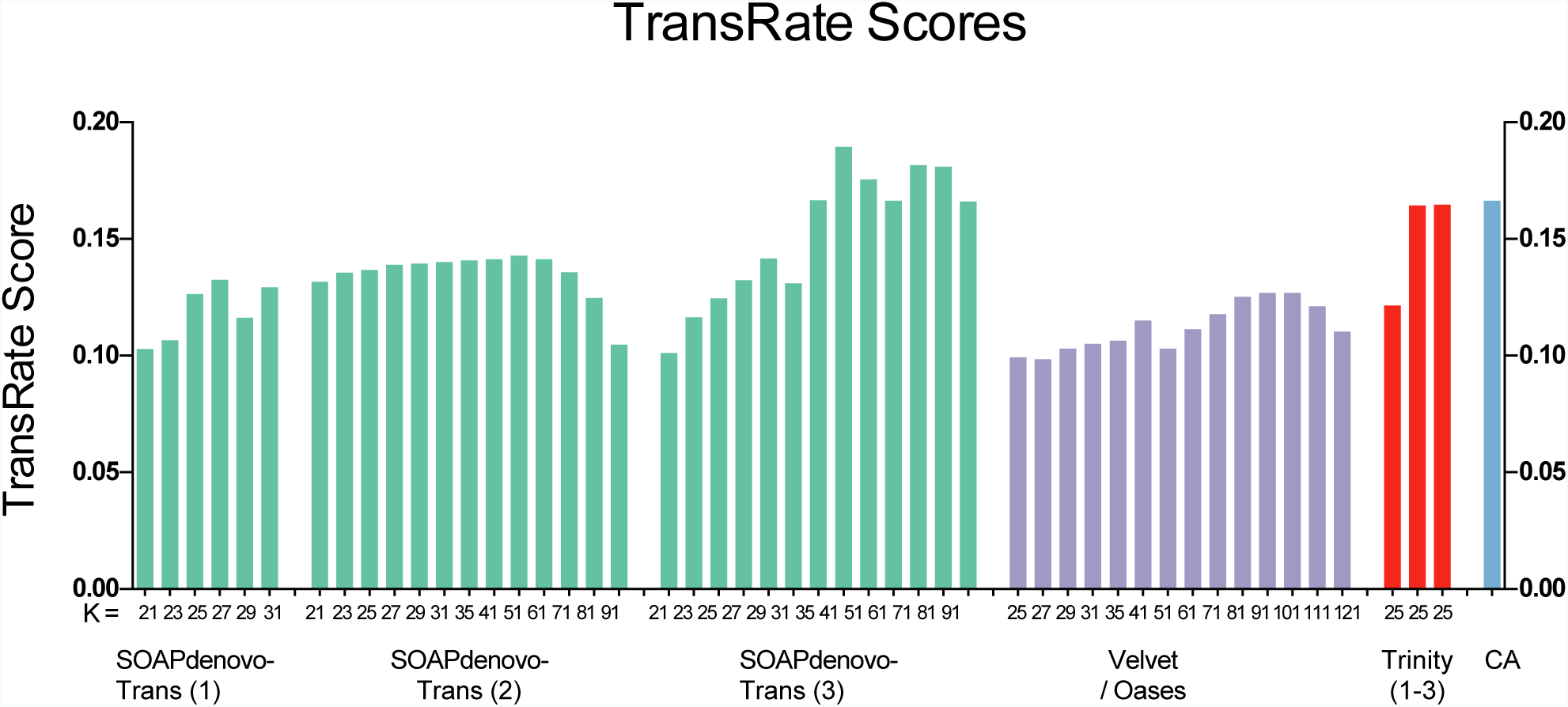
Optimised TransRate scores for each transcriptome assembly. SOAPdenovo-Trans (1): no filtering; SOAPdenovo-Trans (2): light filtering of poor quality read pairs, combined with in silico normalization; SOAPdenovo-Trans (3): moderate filtering of poor quality read pairs; Velvet/Oases: moderate filtering of poor quality read pairs; Trinity (1): light filtering of poor quality read pairs, combined with in silico normalization; Trinity (2): light filtering of poor quality read pairs, combined with SEECER error correction; Trinity (3): moderate filtering of poor quality read pairs; CA: tr2aacds ‘clustered assembly’.

Read errors identified using probabilistic error correction program SEECER were comprised of 14,821,705 substitutions, 1,760,162 deletions, and 1,614,908 insertions, affecting 6% of reads in total. Error correction provided a modest improvement to BUSCO score and mapping of independent reads, however it also resulted in slightly poorer backmapping rate and transrate score, compared to assemblies generated from non-corrected reads (Figures 2 - 5). A recent analysis of RNA-Seq read error correction supports the use of SEECER for enhancing transcriptome assembly quality (MacManes, 2015), however this result suggests the potential benefits of read error correction may not be realized in all *de novo* assembly pipelines.

Trinity produced the most accurate and complete single-kmer assembly comprised of 2,011,167 contigs. Clustering highly similar contigs resulted in a small reduction in the total number of contigs (Supplementary Figure S2), with the majority of clustered contigs found to be gene isoforms (Supplementary Table S3, Table S4, Table S5). Clustering highly similar contigs increased the proportion of single copy BUSCOs detected, however it decreased the proportion of duplicate BUSCOs and increased the number of fragmented BUSCOs (Supplementary Figure S3).

Non-redundant transcripts from all assemblies were merged into a single clustered assembly (CA) using the EvidentialGene *tr2aacds* pipeline (Supplementary Table S4). This was effective in increasing the proportion of complete BUSCOs found, and reducing the number of fragmented and missing BUSCOs (Figure 2). The BUSCO values obtained are consistent with the most complete reference transcriptomes from other vertebrate and eukaryote taxa (Simão et al., 2015: Supplementary Online Material).

### Annotation

The most accurate and complete single-kmer assembly was produced by Trinity from the ‘non-normalized’ dataset. It contained 2,011,167 contigs (1,881,373 ‘genes’ as defined by Trinity), representing a 995 Mb transcriptome. In total, 177,496 transcripts showing significant homology to the Uniprot/Swissprot database, with 59,912 (33.75%) of these transcripts containing an open reading frame, and 7,563 signalling proteins and 1,119 protein families were identified. The output from the annotation pipeline is provided in Supplementary Table S5. In comparison, the *tr2aacds*-generated CA was comprised of 888,080 transcripts, representing a 451 Mb transcriptome, with 189,925 transcripts showed significant homology to the Uniprot/Swissprot database, and with 109,222 (57.5%) containing an open reading frame.

Of the transcripts returning BLASTx hits, 44,706 were associated with one or more Gene Ontology (GO) terms, with 394,288 GO categories represented. GO terms attributed to the greatest number of genes were metal ion binding (GO:0046872; n=13850), RNA-directed DNA polymerase activity (GO:0003964; n=11357), DNA recombination (GO:0006310; n=10728), cytoplasm (GO:0005737; n=9111), nucleus (GO:0005634; n=8648), integral component of membrane (GO:0016021; n=7829), endonuclease activity (GO:0004519; n=7619), plasma membrane (GO:0005886; n=5740), DNA binding (GO:0003677; n=3718), and membrane (GO:0016020; n=3676). GO terms were well distributed between the categories of biological process, cellular component and molecular function (Supplementary Figure S4).

### Validation

Further validation of transcriptome quality was performed by downloading and mapping RNA-Seq reads from NCBI projects: PRJNA184055 and PRJNA292021. A relatively large proportion of reads from these datasets mapped to the transcriptome assemblies generated, with the highest proportion from both datasets mapping to the CA (Figure 3B, Figure 4). Differences in mapping rates between these two independent datasets are likely due to read length (Fushan et al. PRJNA184055: 50 bp paired-end reads; Gawriluk et al. PRJNA292021: 100 bp paired-end reads), and depth of sequencing. Reads used to generate each of the 50 transcriptome assemblies (150bp paired-end reads) were also mapped to each assembly (backmapping), with very few unmapped reads further validating the completeness of the CA (Figure 3). The use of ‘4th generation’ long-read sequencing to improve assembly structure would likely increase the proper pairing of reads, and may reduce the number of unmapped reads further, however the relatively low proportion of unmapped reads is encouraging for broad applications as a reference sequence.

TransRate scores were highest for assemblies generated from non-normalized data, with higher scores typically corresponding to larger kmer sizes (Figure 5). Despite performing well in other measures of transcriptome assembly integrity and completeness, TransRate scores were lower than expected for high-quality transcriptomes (≥0.2), with assemblies typically scoring between 0.1 and 0.2. Possible causes for lower-than-expected TransRate scores are the large proportion of read errors identified by SEECER, and the inclusion of poor quality reads from the original dataset. TransRate scoring is contingent on accurate backmapping of reads to transcripts, as it evaluates assemblies based on whether each base has been called correctly, whether bases are truly part of transcripts, whether contigs are derived from a single transcript, and whether contigs are structurally complete and correct (Smith-Unna et al., 2016). Poor read quality impedes read mapping, which can negatively impact the TransRate score. Aligning error-corrected reads may increase the TransRate score compared to uncorrected reads, however this was not examined here as the format of reads corrected using SEECER is incompatible with TransRate.

## Conclusion

We have generated a comprehensive *de novo* transcriptome assembly for the spiny mouse (*Acomys cahirinus*) using the combined output of three *de novo* transcriptome assemblers: Trinity, SOAPdenovo-Trans, and Velvet/Oases. The EvidentialGene *tr2aacds* pipeline was effective in identifying and collating unique transcripts from 50 unique assemblies, producing a 451 Mb transcriptome assembly comprised of 888,080 transcripts. To our knowledge, this is the first reported instance of the *tr2aacds* pipeline augmenting transcriptome assembly in a mammal. The resulting transcriptome assembly was extensively validated, and shown to be accurate and relatively complete. This transcriptome represents the largest gene catalogue to date for the spiny mouse, providing an important resource for further investigation, and facilitating access to scientific techniques such as sgRNA design for CRISPR/Cas9 gene editing. This reference sequence is now being used to further investigate physiological traits unique to this species, and for establishing the spiny mouse as a useful experimental and clinical model to advance biomedical science.

**Additional file 1: Table S1.** Summary statistics for RNA-Seq runs.

|  | <u>Forward reads</u> |  | <u>Reverse reads</u> |  | <u>Total</u> |
| --- | --- | --- | --- | --- | --- |
| Yield Total (G) | 45.9 |  | 45.9 |  | 91.77617 |
| Aligned (%) | 0.45 |  | 0.25 |  | 0.3509741 |
| Error Rate (%) | 0.62 |  | 1.6 |  | 1.003602 |
| Intensity Cycle 1 | 3255 |  | 3054 |  | 3154.379 |
| % >= Q30 | 80.5 |  | 46.5 |  | 63.47869 |
| Lane | 1 | 2 | 1 | 2 |  |
| Tiles | 64 | 64 | 64 | 64 |  |
| Density (K/mm2) | 1209 ± 78 | 1242 ± 72 | 1209 ± 78 | 1242 ± 72 |  |
| Clusters PF (%) | 64.69 ± 14 | 69.66 ± 10 | 64.69 ± 14 | 69.66 ± 10 |  |
| Phas/Prephas (%) | 0.157 / 0.202 | 0.151 / 0.209 | 0.076 / 0.121 | 0.100 / 0.164 |  |
| Reads (M) | 222.79 | 228.94 | 222.79 | 228.94 |  |
| Reads PF (M) | 145.46 | 160.46 | 145.46 | 160.46 |  |
| % >= Q30 | 80.2 | 80.7 | 42.1 | 50.4 |  |
| Yield(G) | 22 | 24 | 22 | 24 |  |
| Cycles Err Rated | 150 | 150 | 0 - 150 | 0 - 150 |  |
| Aligned (%) | 0.44 ± 0.06 | 0.46 ± 0.05 | 0.22 ± 0.20 | 0.28 ± 0.20 |  |
| Error Rate (%) | 0.65 ± 0.30 | 0.59 ± 0.24 | 1.59 ± 0.51 | 1.61 ± 0.90 |  |
| Error Rate 35 cycle | 0.33 ± 0.26 | 0.24 ± 0.21 | 0.89 ± 0.39 | 0.95 ± 0.61 |  |
| Error Rate 75 cycle | 0.38 ± 0.27 | 0.31 ± 0.22 | 1.27 ± 0.51 | 1.18 ± 0.60 |  |
| Error Rate 100 cycle | 0.44 ± 0.27 | 0.38 ± 0.29 | 1.35 ± 0.52 | 1.30 ± 0.69 |  |
| Intensity Cycle 1 | 3285 ± 266 | 3225 ± 264 | 2995 ± 1829 | 3113 ± 1678 |  |

**Additional file 4: Figure S2.**
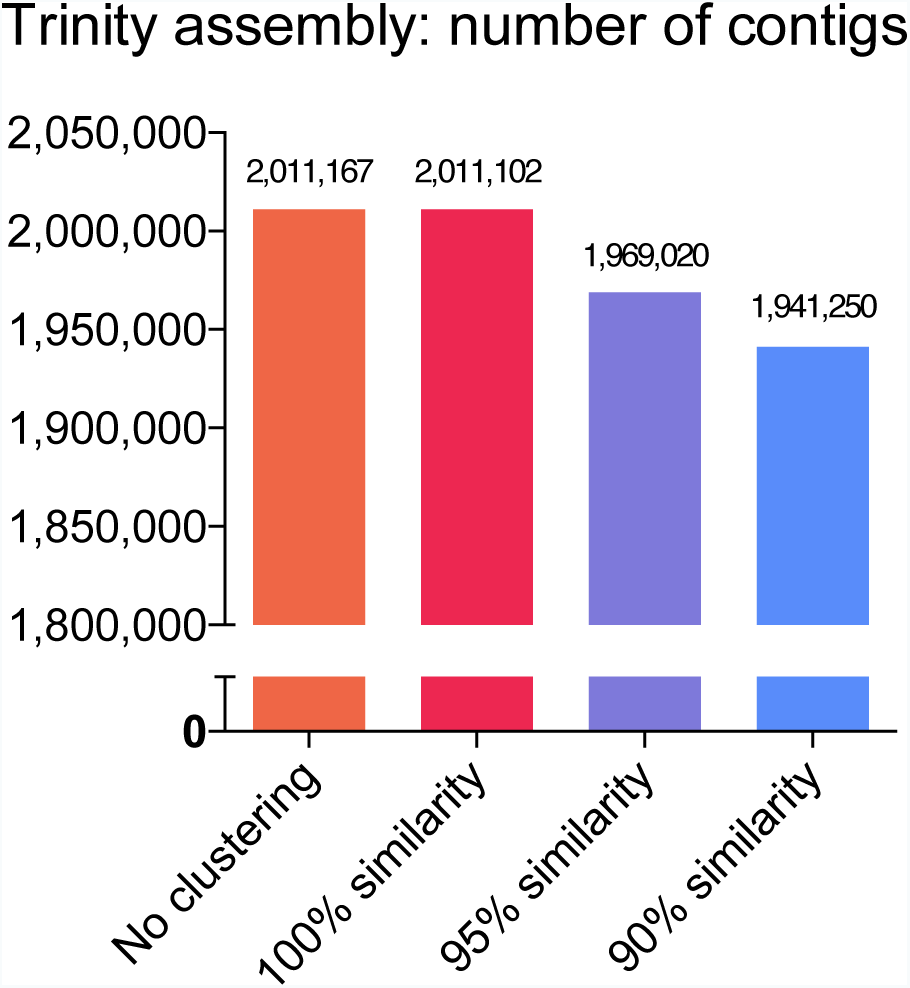
Total number of contigs remaining in Trinity assembly (moderate filtering of poor quality reads, no in silico normalization) after clustering with CD-HIT-EST at each similarity threshold.

**Additional file 5: Figure S3.**
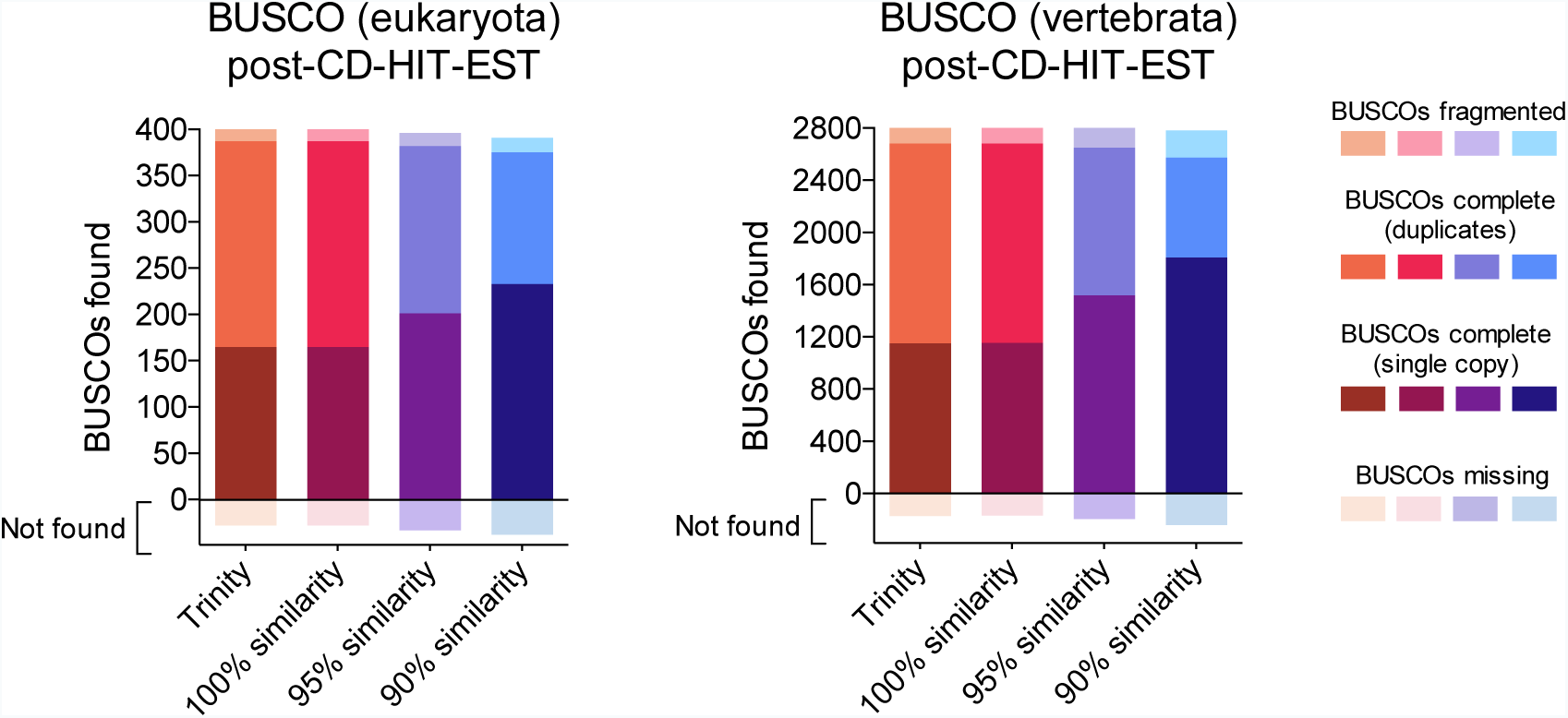
Eukaryote and vertebrate-specific BUSCOs aligning to the Trinity assembly (moderate filtering of poor quality reads, no in silico normalization) before, and after clustering with CD-HIT-EST at specified similarity thresholds

**Additional file 11: Figure S4.**
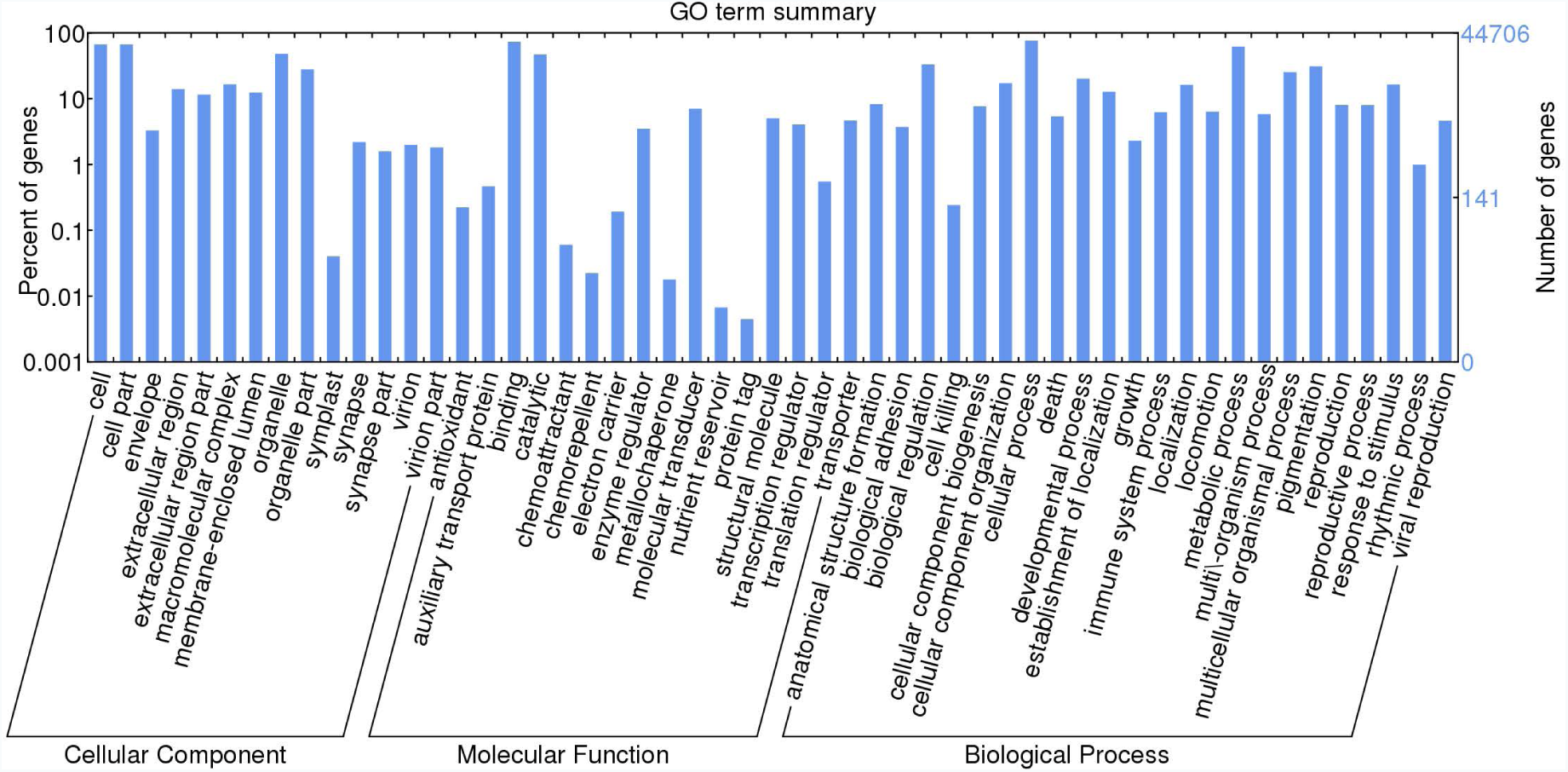
Summary of gene ontology terms (GO terms)

## Abbreviations

bp: base pair
BUSCO: Benchmarking Universal Single-Copy Orthologs
CA: clustered assembly
contig: contiguous sequence
GO: gene ontology
Mb: megabase
NCBI SRA: National Center for Biotechnology Information sequence read archive
NGS: next-generation sequencing
qPCR: quantitative real-time polymerase chain reaction
RNA: ribonucleic acid
RIN: RNA integrity number
RNA-Seq: RNA sequencing
SQL: Structured query language
tr2aacds: transcript to amino acid coding sequence
WEGO: Web Gene Ontology Annotation Plot

## Author contributions

HD and PTS designed the project. HD collected the tissues for sequencing and funded the project. TW conducted the library preparation and sequencing. ATP designed the assembly pipeline. JM conducted the assembly, annotation, analysis and validation, with guidance and advice from ATP and RL. All authors were involved in interpretation of results and all authors read and approved the final manuscript.

## Acknowledgements

The authors would like to thank the following people for their contribution to the study: Daniel Cameron, Gerry Tonkin-Hill and all members of the Papenfuss lab; Torsten Seemann, David Powell, Steve Androulakis and all members of the Monash Bioinformatics Platform; Vivien Vasic and all members of the MHTP Medical Genomics Facility; and Walter and Eliza Hall Institute Department of Bioinformatics.

## References

Athanasiadis, A., Rich, A. and Maas, S. (2004). Widespread A-to-I RNA editing of Alu-containing mRNAs in the human transcriptome. PLoS Biology, 2(12), p.e391. http://dx.doi.org/10.1371/journal.pbio.0020391

Bellofiore, N., Ellery, S.J., Mamrot, J., Walker, D.W., Temple-Smith, P. and Dickinson, H. (2016). First evidence of a menstruating rodent: the spiny mouse (Acomys cahirinus). American Journal of Obstetrics and Gynecology. Aug 5. pii: S0002-9378(16)30476-8. http://dx.doi.org/10.1016/j.ajog.2016.07.041

Bens, M., Sahm, A., Groth, M., Jahn, N., Morhart, M., Holtze, S., Hildebrandt, T.B., Platzer, M. and Szafranski, K. (2016). FRAMA: from RNA-seq data to annotated mRNA assemblies. BMC genomics, 17(1), p.1. http://dx.doi.org/10.1186/s12864-015-2349-8

Bolger, A.M., Lohse, M. and Usadel, B. (2014). Trimmomatic: a flexible trimmer for Illumina sequence data. Bioinformatics, p.btu170. http://dx.doi.org/10.1093/bioinformatics/btu170

Brunjes, P.C. (1990). The precocial mouse, Acomys cahirinus. Psychobiology, 18 (3), pp.339–350. http://dx.doi.org/10.3758/BF03327252

Bushnell, B. (2016). “BBMap short read aligner.” URL http://sourceforge.net/projects/bbmap.

Carneiro, M., Rubin, C.J., Di Palma, F., Albert, F.W., Alföldi, J., Barrio, A.M., Pielberg, G., Rafati, N., Sayyab, S., Turner-Maier, J. and Younis, S. (2014). Rabbit genome analysis reveals a polygenic basis for phenotypic change during domestication. Science, 345(6200), pp.1074–1079. http://dx.doi.org/10.1126/science.1253714

Crusoe, M. R., Alameldin, H. F., Awad, S., et al. (2015). The khmer software package: enabling efficient nucleotide sequence analysis [version 1 referees: 2 approved, 1 approved with reservations]. F1000Research, 4:900. http://dx.doi.org/10.12688/f1000research.6924.1

Dickinson, H., Walker, D.W., Cullen-McEwen, L., Wintour, E.M. and Moritz, K. (2005). The spiny mouse (Acomys cahirinus) completes nephrogenesis before birth. American Journal of Physiology-Renal Physiology, 289(2), pp.F273–F279. http://dx.doi.org/10.1152/ajprenal.00400.2004

Du, L., Li, W., Fan, Z., Shen, F., Yang, M., Wang, Z., Jian, Z., Hou, R., Yue, B. and Zhang, X. (2015). First insights into the giant panda (Ailuropoda melanoleuca) blood transcriptome: a resource for novel gene loci and immunogenetics. Molecular ecology resources, 15(4), pp.1001–1013. http://dx.doi.org/10.1111/1755-0998.12367

Emera, D., Romero, R., and Wagner, G. (2012). The evolution of menstruation: A new model for genetic assimilation. Bioessays, 34 (1), 26–35. http://dx.doi.org/10.1002/bies.201100099

Francischetti, I.M., AssumpÐão, T.C., Ma, D., Li, Y., Vicente, E.C., Uieda, W. and Ribeiro, J.M. (2013). The “Vampirome”: transcriptome and proteome analysis of the principal and accessory submaxillary glands of the vampire bat Desmodus rotundus, a vector of human rabies. Journal of proteomics, 82, pp.288–319. http://dx.doi.org/10.1016/j.jprot.2013.01.009

Fu, L., Niu, B., Zhu, Z., Wu, S., and Li, W. (2012). CD-HIT: accelerated for clustering the next generation sequencing data. Bioinformatics, 28(23), 3150–3152. http://dx.doi.org/10.1093/bioinformatics/bts565

Fushan, A. A., Turanov, A. A., Lee, S. G., Kim, E. B., Lobanov, A. V., Yim, S. H., Buffenstein, R., Lee, S.R., Chang, K.T., Rhee, H. and Kim, J.S. (2015). Gene expression defines natural changes in mammalian lifespan. Aging cell, 14(3), 352–365. http://dx.doi.org/10.1111/acel.12283

Garber, M., Grabherr, M.G., Guttman, M. and Trapnell, C. (2011). Computational methods for transcriptome annotation and quantification using RNA-seq. Nature methods, 8(6), 469–477. http://dx.doi.org/10.1038/nmeth.1613

Gawriluk, T.R., Simkin, J., Thompson, K.L., Biswas, S.K., Clare-Salzler, Z., Kimani, J.M., Kiama, S.G., Smith, J.J., Ezenwa, V.O. and Seifert, A.W. (2016). Comparative analysis of ear-hole closure identifies epimorphic regeneration as a discrete trait in mammals. Nature communications, 7. http://dx.doi.org/10.1038/ncomms11164

Gilbert, D. (2013). EvidentialGene: tr2aacds, mRNA transcript assembly software. URL http://arthropods.eugenes.org/EvidentialGene/

Hadid, Y., Pavlíček, T., Beiles, A., Ianovici, R., Raz, S., and Nevo, E. (2014). Sympatric incipient speciation of spiny mice Acomys at “Evolution Canyon,” Israel. Proceedings of the National Academy of Sciences, 111(3), 1043–1048. http://dx.doi.org/10.1073/pnas.1322301111

Huang, Y., Niu, B., Gao, Y., Fu, L., and Li, W. (2010). CD-HIT Suite: a web server for clustering and comparing biological sequences. Bioinformatics, 26(5), 680–682. http://dx.doi.org/10.1093/bioinformatics/btq003

Lamers, W. H., Mooren, P. G., Griep, H., Endert, E., Degenhart, H. J., and Charles, R. (1986). Hormones in perinatal rat and spiny mouse: relation to altricial and precocial timing of birth. American Journal of Physiology-Endocrinology and Metabolism, 251(1), E78–E85.

Langmead, B., Trapnell, C., Pop, M., and Salzberg, S. L. (2009). Ultrafast and memory-efficient alignment of short DNA sequences to the human genome. Genome biology, 10(3), R25. http://dx.doi.org/10.1186/gb-2009-10-3-r25

Le, H.S., Schulz, M.H., McCauley, B.M., Hinman, V.F. and Bar-Joseph, Z. (2013). Probabilistic error correction for RNA sequencing. Nucleic acids research, p.gkt215. http://dx.doi.org/10.1093/nar/gkt215

Li, W. and Godzik, A. (2006). Cd-hit: a fast program for clustering and comparing large sets of protein or nucleotide sequences. Bioinformatics, 22(13), 1658–1659. http://dx.doi.org/10.1093/bioinformatics/btl158

Li, H., Handsaker, B., Wysoker, A., Fennell, T., Ruan, J., Homer, N., Marth, G., Abecasis, G. and Durbin, R. (2009). The sequence alignment/map format and SAMtools. Bioinformatics, 25(16), pp.2078–2079. http://dx.doi.org/10.1093/bioinformatics/btp352

MacManes, M.D. (2015). Optimizing error correction of RNAseq reads. bioRxiv, p.020123. http://dx.doi.org/10.1101/020123

MacManes, M.D. (2016). Establishing evidenced-based best practice for the de novo assembly and evaluation of transcriptomes from non-model organisms. bioRxiv, p.035642. http://dx.doi.org/10.1101/035642

Martin M. Cutadapt removes adapter sequences from high-throughput sequencing reads. EMBnet journal. 2011;17:10–2. http://dx.doi.org/10.14806/ej.17.1.200

Maudhoo, M.D., Madison, J.D. and Norgren, R.B. (2015). De novo assembly of the chimpanzee transcriptome from NextGen mRNA sequences. GigaScience, 4(1), p.1. http://dx.doi.org/10.1186/s13742-015-0061-x

Nielsen, H., Engelbrecht, J., Brunak, S., and von Heijne, G. (1997). Identification of prokaryotic and eukaryotic signal peptides and prediction of their cleavage sites. Protein engineering, 10(1), 1–6. http://dx.doi.org/10.1093/protein/10.1.1

Patro, R., Duggal, G. Love, M., I., Irizarry, R. A., and Kingsford, C. (2015). Salmon provides accurate, fast, and bias-aware transcript expression estimates using dual-phase inference. bioRxiv, p.021592. http://dx.doi.org/10.1101/021592

Petersen, T. N., Brunak, S., von Heijne, G., and Nielsen, H. (2010). SignalP 4.0: discriminating signal peptides from transmembrane regions. Nature methods, 8(10), 785–786. http://dx.doi.org/10.1038/nmeth.1701

Quinn, T. A., Ratnayake, U., Dickinson, H., Castillo-Melendez, M., and Walker, D. W. (2015). Ontogenetic Change in the Regional Distribution of Dehydroepiandrosterone-Synthesizing Enzyme and the Glucocorticoid Receptor in the Brain of the Spiny Mouse (Acomys cahirinus). Developmental neuroscience, 38(1), 54–73. http://dx.doi.org/10.1159/000438986

Quinn, T. A., Ratnayake, U., Dickinson, H., Nguyen, T. H., McIntosh, M., Castillo-Melendez, M., Conley, A., J., Walker, D. W. (2013). Ontogeny of the adrenal gland in the spiny mouse, with particular reference to production of the steroids cortisol and dehydroepiandrosterone. Endocrinology, 154(3), pp1190–1201. http://dx.doi.org/10.1210/en.2012-1953

Robertson, L.S. and Cornman, R.S. (2014). Transcriptome resources for the frogs Lithobates clamitans and Pseudacris regilla, emphasizing antimicrobial peptides and conserved loci for phylogenetics. Molecular ecology resources,14(1), pp.178–183. http://dx.doi.org/10.1111/1755-0998.12164

Robertson, G., Schein, J., Chiu, R., Corbett, R., Field, M., Jackman, S.D., Mungall, K., Lee, S., Okada, H.M., Qian, J.Q. and Griffith, M. (2010). De novo assembly and analysis of RNA-seq data. Nature methods, 7(11), pp.909–912. http://dx.doi.org/10.1038/nmeth.1517

Schulz, M. H., Zerbino, D. R., Vingron, M., and Birney, E. (2012). Oases: robust de novo RNA-seq assembly across the dynamic range of expression levels. Bioinformatics, 28(8), 1086–1092. http://dx.doi.org/10.1093/bioinformatics/bts094

Shimoyama, M., Smith, J.R., De Pons, J., Tutaj, M., Khampang, P., Hong, W., Erbe, C.B., Ehrlich, G.D., Bakaletz, L.O. and Kerschner, J.E. (2016). The Chinchilla Research Resource Database: resource for an otolaryngology disease model. Database, p.baw073. http://dx.doi.org/10.1093/database/baw073

Simão, F. A., Waterhouse, R. M., Ioannidis, P., Kriventseva, E. V., and Zdobnov, E. M. (2015). BUSCO: assessing genome assembly and annotation completeness with single-copy orthologs. Bioinformatics, 31(19), 3210–3212. http://dx.doi.org/10.1093/bioinformatics/btv351

Smith-Unna, R., Boursnell, C., Patro, R., Hibberd, J. and Kelly, S. (2016). TransRate: reference free quality assessment of de novo transcriptome assemblies. Genome research, pp.gr-196469. http://dx.doi.org/10.1101/gr.196469.115

Surget-Groba, Y. and Montoya-Burgos, J.I. (2010). Optimization of de novo transcriptome assembly from next-generation sequencing data. Genome research, 20(10), pp.1432–1440. http://dx.doi.org/10.1101/gr.103846.109.

Wilson, D.E. and Reeder, D.M. eds. (2005). Mammal species of the world: a taxonomic and geographic reference (Vol. 1). JHU Press.

Xie, Y., Wu, G., Tang, J., Luo, R., Patterson, J., Liu, S., Huang, W., He, G., Gu, S., Li, S. and Zhou, X. (2014). SOAPdenovo-Trans: de novo transcriptome assembly with short RNA-Seq reads. Bioinformatics, 30(12), pp.1660–1666. http://dx.doi.org/10.1093/bioinformatics/btu077

Ye, J., Fang, L., Zheng, H., Zhang, Y., Chen, J., Zhang, Z., Wang, J., Li, S., Li, R., Bolund, L. and Wang, J. (2006). WEGO: a web tool for plotting GO annotations. Nucleic acids research, 34(suppl 2), W293–W297. http://dx.doi.org/10.1093/nar/gkl031

Zaharia, M., Bolosky, W.J., Curtis, K., Fox, A., Patterson, D., Shenker, S., Stoica, I., Karp, R.M. and Sittler, T. (2011). Faster and more accurate sequence alignment with SNAP. ArXiv preprint, arXiv:1111.5572.

Zerbino, D. R., and Birney, E. (2008). Velvet: algorithms for de novo short read assembly using de Bruijn graphs. Genome research, 18(5), 821–829. http://dx.doi.org/10.1101/gr.074492.107

Zieliński, P., Stuglik, M.T., Dudek, K., Konczal, M. and Babik, W. (2014). Development, validation and high-throughput analysis of sequence markers in nonmodel species. Molecular ecology resources, 14(2), pp.352–360. http://dx.doi.org/10.1111/1755-0998.12171

